# Two new threatened species of *Rinorea* (Violaceae), forest trees from East and South Regions, Cameroon

**DOI:** 10.1101/2021.11.05.467439

**Authors:** Gaston Achoundong, Martin Cheek

**Affiliations:** IRAD-Cameroon National Herbarium, BP 1601, Yaoundé, Cameroon; Herbarium, Royal Botanic Gardens, Kew, Richmond, Surrey TW9 3AE, United Kingdom

**Keywords:** Butterflies, conservation, Cymothoe, hydro-electric projects, Lom-Pangar, semi-deciduous forest

## Abstract

Two tree species are described as new to science: *Rinorea spongiocarpa* Achound. sp. nov (placed in *Rinorea* [unranked] Brachypetalae) and *Rinorea dimakoensis* Achound. sp. nov.( placed in *Rinorea* [unranked] Ilicifolieae). Both species are endemic to Cameroon, occurring south of the Sanaga river, the first from South and East Regions, occurring in evergreen forest from Ebolowa to Dja, while the second occurs in the northern part of East Region in semi-deciduous forest towards the interface with woodland habitats in the Dimako-Bertoua area. The two species are illustrated, and their affinities and conservation status according to the 2012 IUCN categories and criteria are discussed. Both species are threatened with extinction due to habitat destruction, the first is assessed as Vulnerable, the second Endangered.

## Introduction

In the course of revising the species of Violaceae of Africa, mainly in preparation for the account of the Violaceae for the “Flore du Cameroun”, the first author has, with collaborators, published 17 new species to science for this group (Achoundong & Onana 1998; Achoundong & Bos 1999; Achoundong & Bos 2001; Achoundong 2003; Achoundong & Cheek 2003; Achoundong & Cheek 2005; Achoundong & Bakker 2006 ; Achoundong *et al.* 2021). The new species published here have previously been referred to under provisional names (Achoundong 1996; Achoundong 1997; Amiet & Achoundong 1996; Achoundong 2000; Bakker *et al.* 2006). In this paper,these two provisional species names are formally published to validate these names.

The most recent studies of the phylogeny and classification of African *Rinorea* Aubl. are set out by Wahlert (2010), Wahlert & Ballard (2012) and van Velzen *et al.* (2015). However, the classification of Brandt (1914) has still not been formally replaced.

The genus *Rinorea* is pantropical, with 212 species currently accepted by Plants of the World Online (POWO, continuously updated, accessed Oct. 2021). Africa is the most species-diverse continent for *Rinorea* with 110 – 150 species (van Velzen *et al.* 2015). *Rinorea* species are mainly forest understorey shrubs or small trees. Morphologically, in continental Africa, they are characterized by having alternate, simple leaves, often with petioles of different lengths on the same stem and a usually long, curving apical bud (in the Neotropics and Madagascar, some species e.g. *Rinorea* sect. *Pubiflora* Wahlert & H. E. Ballard have opposite leaves). The flowers are often green, dull yellow, or shades of white and are zygomorphic. There are three sets of petals in *Rinorea:* an anterior petal (also known as the lower or ventral petal), two lateral petals and two posterior petals. These are likely homologous to the three sets of petals in other strongly zygomorphic genera of Violaceae, such as *Viola* L.(Wahlert 2010).

The anterior petal is larger than the other petals, and often modified, with taxonomically important, often diagnostic characters. The androecium has a staminal tube which is also zygomorphic: the anterior (lower or ventral) side is longer and entire, while on the dorsal side of many of the African species, the staminal tube is generally shorter and incised with a V-shaped cleft. The fruits are typical of the family: hard and dry (except in a few species where it is spongy, see below), tricoccal, septicidal capsules with parietal placentation.

*Rinorea* are ecologically important and diverse in African forests, often with several sympatric species in one forest. For example, ten species were recorded in a few square kilometers of the Mefou Proposed National Park near Yaoundé (Achoundong in Cheek *et al.* 2011). Many species are range-restricted, found in such small areas that they are at risk of extinction from forest clearance. *Rinorea dewitii* Achound., *R. fausteana* Achound., *R. simoneae* Achound. and *R. thomasii* Achound. are all assessed as threatened in the Red Data Book of the Flowering Plants of Cameroon (Onana & Cheek 2011) and all but the first can be found on the IUCN Red List (https://iucnredlist.org) e.g., *Rinorea thomasii* (Darbyshire & Cheek 2004a; Cheek 2017; Darbyshire & Cheek 2004b). In neighbouring Gabon, the recently published *Rinorea calcicola* Velzen & Wieringa is also range-restricted and of conservation concern (van Velzen & Wieringa 2014). Cameroon has the highest species-diversity for the genus in tropical Africa with 53 species listed (Onana 2011), followed by Gabon, with 46 species (Sosef *et al.* 2006). However, the superficial similarity between species has made identification difficult for taxonomists, e.g., 194 specimens of *Rinorea* are listed as unidentified to species for Gabon in Sosef *et al.* (2006).

African *Rinorea* species are of great interest to entomologists, being important larval food plants of the butterfly genus *Cymothoe* (the gliders). Twenty-eight species of *Cymothoe* are known to feed on *Rinorea*, of which 18 are strictly monophagous, six are oligophagous and three feed on up to six species of *Rinorea* (Amiet 1997; Amiet 2000; Amiet & Achoundong, 1996).

## Materials & Methods

Fieldwork by the first author was mainly carried out in Cameroon from1987 – 1996 in connection with his doctoral studies of *Rinorea* (Achoundong 1997). All specimens cited have been seen. Herbarium citations follow Index Herbariorum (Thiers *et al.* continuously updated). Specimens were studied online, on loan from or at BR, K, P, WAG and YA principally by the first author. We also searched JSTOR Global Plants (https://plants.jstor.org/ accessed April 2021) for additional materials. Taxonomic authorities follow the International Plant Names Index (IPNI 2021), and nomenclature follows Turland *et al.* (2018). The conservation assessment was made using Bachman *et al.* (2011) following the categories and criteria of IUCN (2012). Herbarium material was examined with a Leica Wild M8 dissecting binocular microscope fitted with an eyepiece graticule. Measurements were made from rehydrated or spirit preserved material. The terms and format of the description follow the conventions of Achoundong & Cheek (2005) and Achoundong *et al.* (2021). Post-facto georeferences for specimens without coordinates were obtained from Google Earth. (https://www.google.com/intl/en_uk/earth/versions/).

## Results: Taxonomy

*Rinorea spongiocarpa* Achound. sp. nov., the first of the two species described below is placed in *Rinorea* [unranked] Brachypetalae because it is closely similar to *R. gabunensis* and *R. leiophylla* which are placed in this group on molecular grounds (Wahlert and Ballard 2012; van Welzen *et al.* 2015) and because it fits the description of this group: alternate leaves, six ovules per ovary, cymose inflorescence, anthers sessile on the border of the staminal tube.

*Rinorea dimakoensis* Achound. sp. nov. the second species described, is placed in *Rinorea* [unranked] Ilicifolieae on molecular grounds (Wahlert and Ballard 2012; van Welzen *et al.* 2015) and because it fits the description of the group: alternate leaves, six ovules per ovary ; cymose inflorescences, sepals clearly ribbed fanwise; staminal tube sinuate between insertion of anthers, tube without a free margin or a lobed margin subtending anthers. Further data are given on Ilicifoliae in Wahlert *et al.* (2020).

### 1. Rinorea spongiocarpa Achound

**sp nov** Type. Cameroon, South Region, Ebolowa, “Hill facing village of N’Kolandom hill, in primary forest on slope, Alt. c.700 m, 2.48N, 11.10E”, fl. 20 Feb.1975, *J.J*. *F.E. de Wilde* 7985 (holotype WAG; isotypes BR, K000593339, P, YA). (Fig. 1).

**Fig. 1.**
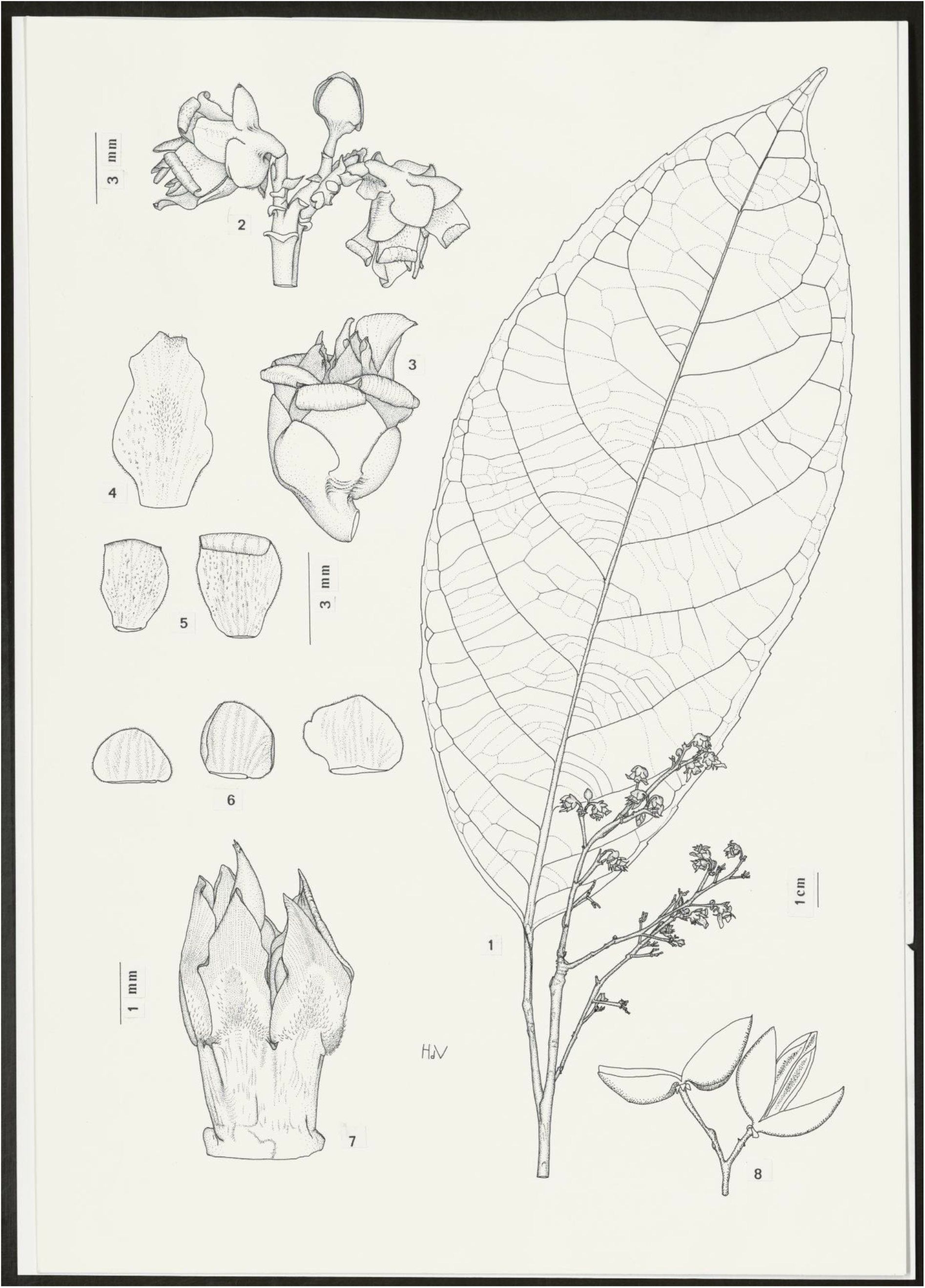
Rinorea spongiocarpa. **A** habit, flowering stem; **B** portion of inflorescence with two open flowers and one flower in bud; **C** flower, note the spreading anterior petal; **D** anterior petals; **E** lateral and posterior petals; **F** sepals; **G** androecium, side view; **H** fruit, mature. All drawn from *J.J*. *F.E. de Wilde* 7798 (holotype, WAG) by J.M. (HANS) DE VRIES

**Fig. 2.**
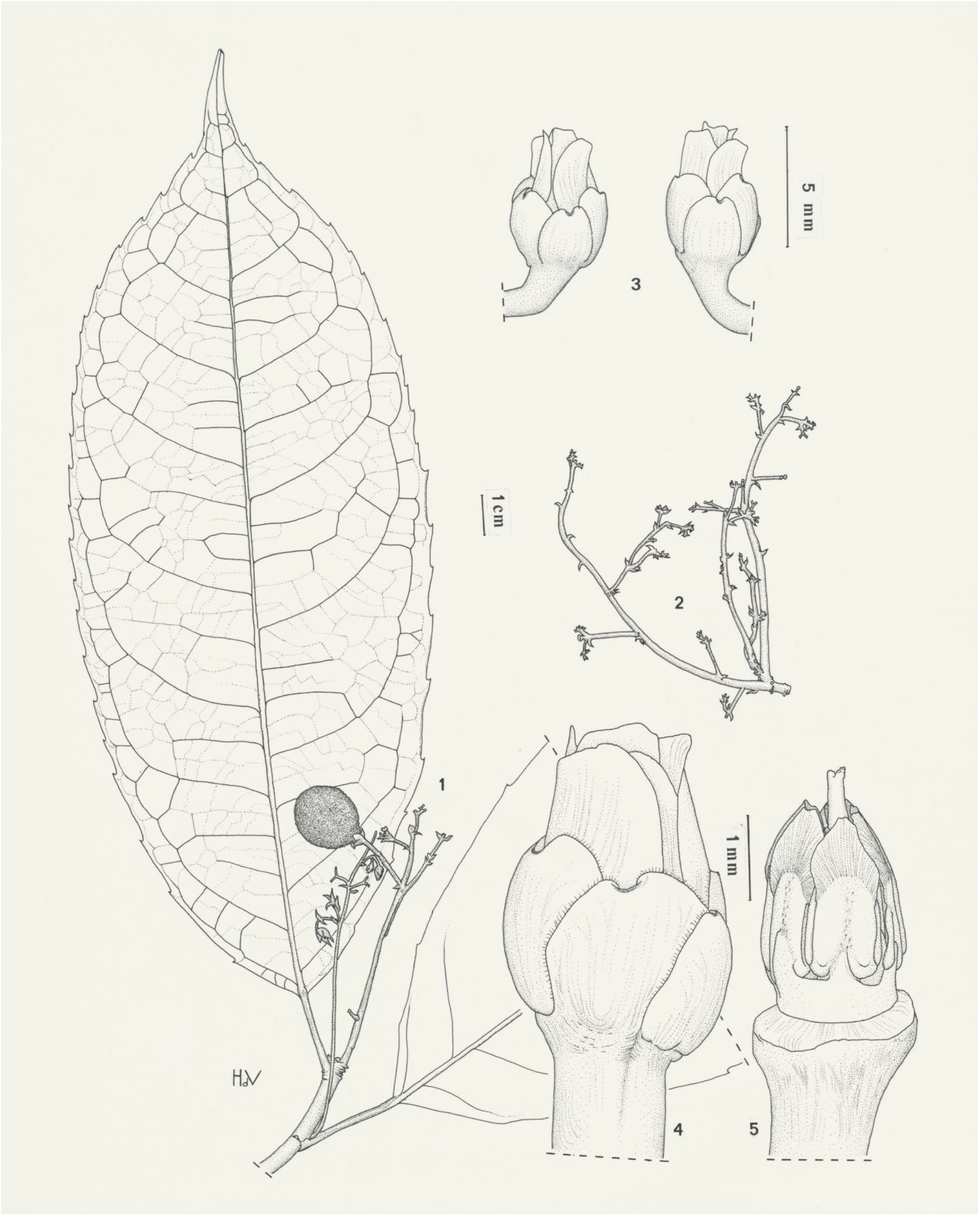
Rinorea dimakoensis. **A** habit, fruiting stem; **B** portion of infructescence axis; **C** two flowers, side view; **D** flower, side view; **E** sepals; **F** androecium, side view; **G** fruit, mature. **A, C-D** from *Achoundong 3033* (WAG, YA), **B** from *Letouzey* 2663 (holotype P). Drawn by J.M.(HANS) DE VRIES

*Rinorea spongiocarpa* ined. Achoundong (1996: 536 – 544; 1997: 193 – 198); Amiet & Achoundong (1996: 465); Onana (2011: 151).

*Tree or shrub* 2 – 6( – 12) m high; stems glabrous. *Leaf-blades* leathery to thickly papery, glossy, dark green above, pale yellow-green beneath, elliptic, ovate to narrowly oblong, 14 – 30 × 11 – 6 cm, apex acuminate, base cuneate to attenuate, lateral nerves 9 – 12, on each side of the midrib, tertiary nerves subscalariform, leaf margin crenate, glabrous; petiole 5 – 7 cm long. *Inflorescence* a terminal panicle up to 7 cm long, lateral ramifications constituted by few-flowered cymes of 5 – 7 flowers each. Bracts triangular, 1.5 × 2 mm, median nerve raised, conspicuous. *Flowers* yellow or yellow-white, zygomorphic, 4 – 5 ( – 6) × 3 – 4 ( – 5) mm. *Sepals* purple, unequal, triangular to elliptic 2 × 3 mm, apex rounded. *Petals* yellow, unequal, anterior (lower) petal oblong 6 × 2 mm, spreading, not or barely revolute at maturity, lateral and posterior petals smaller, 3 – 4 mm long, distal half strongly revolute. *Androecium* zygomorphic, 3.5 – 5 mm long, staminal tube 1 – 2.5 mm long, tube margin not free, anthers sessile on the tube border, thecae 2 mm long, outer surface puberulent, connective appendage 2 mm long, red, decurrent deeply on the anther thecae, thecae appendage entire, not bifid. *Gynoecium* up to 5.3 mm long. Ovary subglobose, 1.5 × 1 mm, glabrous, style straight, enlarged at the base, 3.5 mm long. *Fruit* ovoid, 30 × 20 mm surface smooth, lacking ribs, fruit wall c. 3 mm thick, spongy, six-seeded. *Seeds* white, tetrahedric, 9 – 10( – 11) × 9 – 10 × 5( – 6) mm, drying pale brown, glossy.

#### RECOGNITION

Similar to *R. parviflora* Chipp in the fruits having a thick spongy mesocarp (in almost all other members of the genus the mesocarp is slender and hard), differing in that the abaxial surface of the leaf-blade lacks glands and that the inflorescence branches are many-flowered (vs glandular and 1 – 2-flowered respectively).

#### DISTRIBUTION

Cameroon. The species is restricted to the area of the Dja Forest, East Region, extending westwards to Ngovayang and Ebolowa in Littoral Region.

#### SPECIMENS STUDIED. CAMEROON. East Region

Alat Makay, Dja National Park, fl.23 Feb.1987, *Achoundong* 1411 (P, YA); Timbe II, near Abong-Mbang, fl. 20 Jan.1990, *Achoundong* 1562 (P, YA); Dja National Park., fl. 25 Apr.1993, *Lejoly & Sonké* 154 (BR, YA); Dja National Park, Mekas, fl. 10 Jan.1995, *Sonké* 1385 (BR, YA); Mekas, fr. 26 May 1995, fr, *Sonké* 1548 (BR, YA); **South Region**. Ebolowa, Hill facing village of N’Kolandom hill, in primary forest on slope, fl. 20 Feb. 1975, *J.J*. *F.E. de Wilde* 7985 (holotype WAG; isotypes BR, K000593339, P, YA); Nkoladom village, south of Ebolowa, fr. 10 Sept. 1989, *Achoundong* 1495, (K000593338, WAG, YA); ibid., fl. fr. Sept. 1992, *Achoundong* 1951 (YA); ibid. *Achoundong* 1952 (YA); ibid. *Achoundong* 1953 (YA); Bongolo I, 30 km on Ebolowa-Lolodorf road, fl., 14 Sept. 1989, *Achoundong* 1501, (K, YA); ibid., fl., fr., 14 Sept. 1989, *Achoundong* 1501 (K000593337, P, YA); Avobengon village, 24 km south of Djoum, 12° 55’E, 2° 40’E, fl., 22 Dec. 1990, *Achoundong* 1598 (YA); ibid. fl., 22 Dec.1990, *Achoundong* 1632 (YA); ibid. fl. 22 Dec.1990, *Achoundong* 1700 (P, YA) ; ibid., fl. 10 April 1991, *Achoundong* 1760 (K000593336, WAG, YA); ibid. imm.fr. *Achoundong* 1802 (K000593335, YA); Medjap, near Djoum, st. 20 May 1990, *Achoundong* 1681 (P, YA); Mill Hill, Lolodorf, Sept. 1992, *Achoundong* 1974 (YA); Bibondi near Ngovayang, north West of Lolodorf, fl., 4 March 1993, *Achoundong* 2017 (YA); ibid. *Achoundong* 2024 (YA); Mezese, fr. 24 May1993, *Achoundong* 2066, (YA); Mezese, fr. 16 Sept. 2004, *Achoundong* 2335 (YA, WAG); Ebienemeyong, fl., Jan.1993, *Achoundong* 2338 (YA).

#### HABITAT

This species is widespread in dense lowland evergreen forest of the Cameroon Congolese forest zone (in the sense of Letouzey 1985). It occurs at an altitudinal range of 400-700 m. It does not occur in the littoral plain and is completely absent from the semi-deciduous forest of the South Cameroon Plateau.

#### CONSERVATION STATUS

*Rinorea spongiocarpa* is only known from parts of Littoral Region and the Dja Forest area of adjoining East Region. On the basis of the specimen records cited above, we calculate the total extent of occurrence of *Rinorea spongiocarpa* as 25,295 km^2^. However, within this fairly large area, there are currently only nine scattered locations known and the global area of occupation is calculated as only 44 km^2^ using the IUCN-preferred cell size of 4 km^2^. Surveys for plant conservation management in forest areas north, south and west of the range of distribution of this species have resulted in many thousands of specimens being collected and identified, but failed to find any additional specimens of *Rinorea spongiocarpa* (Cheek 1992 ; Cable & Cheek 1998; Cheek *et al.* 2000; Maisels *et al.* 2000 ; Chapman & Chapman 2001; Tchoutou 2004; Harvey *et al.* 2004; Cheek *et al.* 2004; Cheek *et al.* 2010; Harvey *et al.* 2010; Cheek *et al.* 2011). Although there are still poorly sampled forest locations with intact natural habitat in Cameroon, it is likely that the known range of *Rinorea spongiocarpa* is close to the actual. Only one location for the species occurs in a designated protected area: the Dja forest. The species appears absent from the largest protected area in South Region, the Campo-Ma’an National Park. Despite being formally unprotected, forest at several of the known locations appears mainly intact (Google Earth imagery viewed 29 Oct. 2021), with forest entirely or largely removed only in two or three locations, apparently due to small-holder agricultural operations near roads. *Rinorea spongiocarpa* is therefore here assessed, on the basis of the ten or less locations, small known area of occupation and threats described above, as Vulnerable, VU B2ab(iii). Another example of a range-restricted endemic species in the same range and habitat is *Kupeantha spathulata* (A.P.Davis & Sonké)Cheek (Cheek *et al.* 2018a), which is also assessed as Vulnerable (Rokni *et al.* 2017).

#### ETYMOLOGY

The name of this species derives from its fruit which has a well-developed, thick, spongy mesocarp, while other most species of the genus have slender, more hard-walled fruits.

#### NOTES

In herbarium collections, specimens belonging to this species appear similar to those of two other species: *Rinorea gabunensis* Engl. and *Rinorea leiophylla* M.Brandt to which *Rinorea spongiocarpa* has long been confused. The species have separate ranges. *Rinorea gabunensis* and *Rinorea leiophylla* occur exclusively in the littoral plain whereas *Rinorea spongicarpa* is restricted to forest inland in the Cameroon “congolese forest” zone. The three species are distinguished as follows:

1. Small shrub of less than 5 m high; lamina with secondary nerves prominent; partial-inflorescences (secondary branches) of the inflorescence up to 12 mm long; fruit ribbed………………………………………………………*Rinorea gabunensis*
1. Small tree up to 12 m high; lamina secondary nerves not prominent; partial-inflorescences (secondary branches) of the inflorescence less than 1.4 mm long; fruit smooth (not ribbed)………………………………………………………2
2. Leaves glandular on abaxial surface; staminal tube with free margin……………………………………………………….*Rinorea leiophylla*
2. Leaves not glandular on abaxial surface; staminal tube margin not free………………………………………………………Rinorea spongiocarpa

### 2. Rinorea dimakoensis

Achound. sp. nov. Type. Cameroon, East Region, Dimako Bonda river, fl. fr.18 Jan.1960, *Letouzey* 2663 (holotype P; isotype YA)

*Rinorea dimakoensis* Achound. ined. Achoundong (1996 : 544); (1997 : 156); Amiet & Achoundong (1996: 466); Bakker *et al.* (2006); Onana (2011); van Velzen *et al.* (2015); Wahlert (2010); Wahlert & Ballard (2012).

*Tree or shrub* up to 12 m tall; stems glabrous. *Leaves* thickly coriaceous, dark green above, blades elliptic, ovate to narrowly oblong, 30 – 10 × 6 – 18 cm, apex acuminate, base rounded, obtuse or attenuate, lateral nerves 8 – 10 on each side of the midrib, leaf margin acutely serrate, glabrous; petiole 3 – 6 cm long. *Inflorescence* terminal or subterminal panicle 8 – 12 × 2 – 7 cm, lateral branches with 2 – 5-flowered cymes; bracts ovate 1.5 – 2 × 1 – 2 mm, rounded at the summit, median nerve conspicuous. *Flower* yellow, zygomorphic, 4 – 5 ( – 6) × 3 – 4( – 5) mm. *Sepals* unequal, triangular to elliptic orbicular, 2 × 3 mm, apex rounded or emarginate. *Petals* yellow, unequal, oblong, 6 – 7 × 2 – 4.2 mm, lateral and dorsal petals smaller, anterior (lower) petal bigger, straight, not revolute at maturity. *Androecium* zygomorphic, 3.5 – 5 mm long. Staminal tube 0.5 mm long, tube margin not free, anthers sessile on the rim of the staminal tube; staminal thecae 2 mm long, connective subelliptic, c. 2 mm long, apex rounded, thecae base decurrent slightly, thecal appendage entire (not bifid). *Gynoecium* 3 mm long. Ovary glabrous, subglobose, 1 mm long, style 2 mm long. *Fruit* ovoid, 3.5 × 3 cm, longitudinally 3-ribbed, 6-seeded. Seeds tetrahedric, 8 × 5 mm.

#### RECOGNITION

*Rinorea dimakoensis* Achound. is similar to *R. ilicifolia* Kuntze, in the shape and size of the leaves which are leathery and robustly toothed at the margin, however in *Rinorea dimakoensis* the leaves are wider ( 6 – 18 cm vs 3 – 9 cm) and the spines are shorter (<0.5 mm long vs 1 – 2 mm long), the inflorescences, pedicels, flowers and fruits are longer and/or larger, reaching 8 – 12 cm, 4.5 mm, 7 × 4 mm, and 3.5 cm long respectively (vs 3 – 5 cm, 1 mm, 3.5 – 4 × 3 – 4 mm, and 1.2 – 1.6 cm long and the sepals have only the midrib raised and partly rib-like, not with multiple raised longitudinal ribs.

#### DISTRIBUTION

Cameroon. The species is globally endemic to a narrow area on the South Cameroon Plateau of East Region, extending from Dimako in the south to the Lom-Pangar in the north.

#### SPECIMENS STUDIED. CAMEROON. East Region

South Dimako, Bonda River, fl.fr.10 Aug.1987, *Achoundong* 1878 (P, YA); Confluence du Lom et du Pangar, fl. date unknown, *Achoundong* 3033 (WAG, YA); South of Dimako, Bonda river, fl. fr. 18 Jan. 1960, *Letouzey* 2663 (holotype P, isotype YA); Ndemb II, 55 km along Ndemba road, st. 17 Nov. 1955, *Nana* 343 (P, YA); 60 km on Bertoua Road., Essengue ll path, st. 18 Feb.1956, *Nana* 489 (YA).

### HABITAT

*Rinorea dimakoensis* is globally restricted to semi-deciduous forest at 680 – 720 m alt. The distribution range of this species falls within the transition from forest to grassland in Cameroon, with forest mainly along drainage lines interdigitating with grassland on better-drained areas. The species is completely absent from semi-deciduous forest in the adjoining Centre Region, e.g in the Yaounde and Bafia areas. It is also completely absent from Cameroon coastal (littoral) evergreen forest, where its relative *Rinorea ilicifolia* occurs.

### CONSERVATION STATUS

Formerly three locations were known for *Rinorea dimakoensis.* However, the species has been lost at its former northernmost location which has now been inundated (viewed on Google Earth 29 0ct. 2021) by the reservoir behind the Lom-Pangar hydro-electric dam, which was completed in 2017. It is to be hoped that searching in surviving suitable habitat in the area might discover additional individuals but this is not certain. Forest at the remaining two locations appears to have been shrinking and has been degraded over recent years, probably due to urbanisation, and the demand for fuel in neighbouring Dimako and Bertoua, towns along the transnational highway that links Douala, Yaounde and Bangui, the major artery for the Central African Republic. On the basis of the specimen records cited above, we estimate the total extent of occurrence of *Rinorea dimakoensis* as 705 km2 (including the Lom-Pangar site) and the global area of occupation is calculated as only 8 km^2^ using the IUCN-preferred cell size of 4 km^2^. There are currently only two extant locations known. It is possible that additional locations will be found within the extent of occcurence and that this itself might be extended since this area of Cameroon is less well surveyed than that of the main forest zone (see references cited under *Rinorea spongiocarpa*). However, while it is possible that additional locations might be found, it may also be that the species is truly as localised, range-restricted and threatened as the data indicates, as are several other species of the genus in Cameroon (see introduction above). We here assess *Rinorea dimakoensis* on the basis of the data given above, as Endangered, EN B1+B2ab(iii). Another example of a range-restricted endemic species in the same range is *Alliphylus bertoua* Cheek, also Endangered (Cheek & Haba 2016).

### ETYMOLOGY

The species is named after the Dimako locality in East Region, where the first fertile specimen representing this species was collected in 1960 by Letouzey.

### NOTES

*Rinorea dimakoensis* appears both closely similar and related to *R. ilicifolia* Engl. From field observations it appears that *Rinorea dimakoensis* differs in the shape and size of the habit, the structure of the leaf margin, the architecture of the inflorescence and the habitat. This close similarity probably constitutes one of the reasons why this species has not been recognized previously. In fact the first specimen of this species collected by Letouzey was erroneously identified as *Rinorea ilicifolia*. Since discriminating characters are mainly located in mature flowers, it was not easy for botanists formerly to separate this species. *Rinorea dimakoensis* is also similar to *Rinorea keayii*. However, in *Rinorea dimakoensis* leaves margins are so not spiny as those of this species and the lamina is not glandular beneath.

## Discussion

Documented global extinctions of plant species are increasing (Humphreys *et al.* 2019) and recent estimates suggest that as many as two fifths of the world’s plant species are now threatened with extinction (Nic Lughadha *et al.* 2020). Cameroon has the highest documented number of plant species extinctions of any country in tropical Africa (Humphreys *et al.* 2019). The endemic Cameroon species *Oxygyne triandra* Schltr. and *Afrothismia pachyantha* Schltr. are among those now known to be globally extinct (Cheek & Williams 1999, Cheek *et al.* 2018b, Cheek *et al.* 2019) and two recently two species of *Pseudohydrosme* (Moxon-Holt & Cheek 2021; Cheek *et al.* 2021) have been shown to be extinct in adjoining Gabon. In some cases, species appear to have become extinct even before they are known to science, such as *Vepris bali* Cheek (Cheek *et al.* 2018c), also in Cameroon. Even areas known to be of high conservation value have been slated for development, threatening the species they contain with extinction, e.g. the Ebo forest in Cameroon (Lovell 2020).

About 2000 plant species new to science are published each year, with Cameroon contributing more than any other tropical African country in 2019 (Cheek *et al.* 2020). Only when such species as *Rinorea spongiocarpa* and *R. dimakoensis* (this paper) are formally known to science are they fully visible and can extinction risk assessments be accepted by IUCN allowing the possibility of measures being taken to protect them (Cheek *et al.* 2020).

Efforts are now being made to delimit the highest priority areas in Cameroon for plant conservation as Tropical Important Plant Areas (TIPAs) using the revised IPA criteria set out in Darbyshire *et al.* (2017). This is expected to help avoid the global extinction of additional endemic species such as the *Rinorea* species published in this paper which it is intended will be included in the future TIPAs.

## Acknowledgements

The first author thanks IRAD-National Herbarium of Cameroon (YA) for support in his retirement that has enabled him to continue and finalise his taxonomic research on Violaceae of Cameroon for the Flore Du Cameroun. The Nederlandese Organisatie voor Wetenschappelyk Onderzoek (NOW), Institut de Recherche pour le Developement (IRD) and Royal Botanic Gardens, Kew have supported the cost of visits by the first author to the European herbaria of BR, BM, K, P and WAG. The first author is particularly grateful to Jos van der Maesen, Frans Breteler and Eric Chenin at these institutions. He also thanks colleagues at Wageningen for retrieving and transmitting the excellent figures that illustrate this paper, drawn by J.M. (Hans) de Vries. This paper was completed as part of the Cameroon TIPAs (Tropical Important Plant Areas) project at RBG, Kew, which is supported by players of People’s Postcode Lottery. Formal Red Listing of these two tree species, once they are formally published, will be supported by the John S. Cohen Foundation.

